# Exploration of individual beta cell function over time *in vivo:* effects of hyperglycemia and glucagon-like peptide-1 receptor (GLP1R) agonism

**DOI:** 10.1101/2025.03.31.646461

**Authors:** Luis Fernando Delgadillo-Silva, Shadai Salazar, Livia Lopez Noriega, Audrey Provencher-Girard, Sandra Larouche, Alexandre Prat, Guy A. Rutter

## Abstract

The coordinated function of beta cells within the pancreatic islet is required for the normal regulation of insulin secretion and is partly controlled by specialized “leader” and highly connected “hub” beta-cell subpopulations. Whether cells within these subpopulations are functionally stable *in vivo* remains unclear. Here, we establish an approach to monitor Ca^2+^ dynamics within individual beta cells over time, after engraftment into the anterior eye chamber, where continuous blood perfusion and near normal innervation pertain. Under normoglycemic conditions, islet network dynamics, and the behavior of individual leaders and hubs, remain stable for at least seven days. Hyperglycemia, resulting from high-fat diet feeding or the loss of a host *Gck* allele, caused engrafted islets to display incomplete and abortive Ca^2+^ waves and overall connectivity was diminished. Whereas hub cell numbers were lowered profoundly in both disease models, leaders largely persisted. Treatment with the GLP1R agonist Exendin-4 led to a recovery of islet-wide Ca^2+^ dynamics and the re-emergence of hub cells within minutes, with the effects of the incretin mimetic being more marked than those observed after analogous treatments in *vitro*. Similar observations were made using 3-dimensional imaging across the whole islet. Our findings thus suggest that incretins may act both directly and indirectly on beta cells in vivo. The approach described may provide broad applicability to the exploration of individual cell function over time in the living animal.

## Introduction

All forms of diabetes mellitus are characterized by defective insulin secretion. More than 1 in 10 of the adult population worldwide is affected by diabetes (https://www.idf.org/). Whereas autoimmune destruction of beta cells is usually the cause of type 1 diabetes ^1^, the more prevalent type 2 diabetes (T2D) is caused by beta cell dysfunction and insulin resistance. T2D is driven by both genetic and environmental factors, the latter notably including obesity which interferes with insulin secretion and action ^2^. People with T2D usually have impairments in both phases of insulin secretion ^3,4^ and oscillatory secretion is lost in the relatives of patients with T2D ^5^. Hence, understanding the mechanisms responsible for coordinated insulin secretion is important to develop effective therapies for diabetes.

The mechanisms through which beta cells respond to a rise in blood glucose concentration have been explored in detail ^6^. In brief, these involve enhanced flux through the glucose phosphorylating enzyme, glucokinase, accelerated mitochondrial metabolism and ATP synthesis. Closure of ATP-sensitive K^+^ channels ^7^ then leads to Ca^2+^ influx and the fusion of insulin-containing granules at the cell membrane. Other mechanisms, including locally high levels of glycolytic intermediates, may also be involved ^8^, though their role is disputed ^9^. Incretin hormones, including glucagon-like peptide 1 (GLP-1), potentiate the effects of glucose largely via increases in intracellular cAMP ^10^. In the past, beta cells were often considered as a homogeneous population, forming a uniform functional syncytium ^11^. However, more recent studies have demonstrated that these cells display marked heterogeneity in their insulin-secreting capabilities and sensitivity to glucose ^12-14^, the presence of cell-surface markers ^15^, gene expression profiles ^16,17^ and developmental history ^18,19^. Furthermore, we ^20-23^ and others ^24-26^ have shown that functionally discrete subpopulations of beta cells within the intact isolated islet initiate and coordinate overall Ca^2+^ responses to glucose (e.g. “Leaders” ^21,22^), which initiate Ca^2+^ waves, “hubs” ^20^ connecting the islet-wide network, and “first-responders” ^23,25^ which are involved in the first phase response. The characterization of discrete functional beta cell subpopulations has become the focus of intense investigation, as recently reviewed ^27^. However, most of our understanding of beta cell subpopulations has emerged from single time-point observations. Hence, it remains unknown if these subpopulations represent transient “states” or if they belong to stable, long-lived functional groups ^28^.

In an earlier study ^29^ we reported, using islet imaging in the anterior eye chamber (ACE), changes in islet connectivity during metabolic (high fat) stress, and its recovery after bariatric surgery. However, these studies did not provide a means of exploring the stability of *individual* beta cell subgroups (leaders, hubs, followers) over time. Moreover, only diet-induced metabolic stress was explored, while the effects of hyperglycemia prompted by genetic causes were not examined.

Here, we establish an approach through which the stability of individual leader and hub beta cells can be explored longitudinally *in vivo* over days to weeks, and explore their responses to metabolic challenges provoked by diet or genetic changes, and to acute GLP1 receptor (GLP1R) agonism. Notably, we deploy the cryptic photoprotein PA-mCherry under conditions which allow us to generate stable “landmarks” across the islet (“spontaneous photolabelling”) ^22^. Using this approach, we report the stability of beta cell populations with cellular resolution, the impact of mild dysglycemia, and recovery in response to incretin receptor agonism.

## Methods

### Animal husbandry

All experiments were performed with ethical permission from the CRCHUM Animal Facility (CIPA 2022–10,040 CM21022GRs). Colonies of Ins1Cre:GCaMP6^f/f^ and Gck-mCardinal (J-Gck^tm1(mCard)/Rutt^, Gck^*KI/+*^) ^30^ mice maintained on a C57BL/6J background, were fed a regular chow diet. Ins1Cre:GCaMP6^f/f^ mice express the Ca^2+^ sensor GCaMP6f selectively in the beta cell ^21^ and were used as donors. C57BL/6J wild type (WT; 18-28g) mice were purchased from the Jackson laboratory (Jax.org). Both Gck^*KI/+*^ and purchased WT mice were used as donor-islet recipients. WT mice were fed either a regular chow or High-fat-high-sucrose (HFHS) diet (Cederlane, 58% Fat, D17082304). All groups were maintained under conditions of controlled temperature (21–23 °C), humidity (45–50 %) and light (12 h day-night cycle).

### Islet transplantation into mouse anterior chamber of the eye (ACE)

Pancreatic islets were isolated and cultured as described previously ^31^. For transplantation, 20-30 islets were aspirated with a 27-gauge blunt eye cannula (Labtician, Can) connected to a 0.4-mm polyethylene tubing (Portex Limited) and to a 100ul Hamilton syringe (Hamilton). Mice were provided with analgesic Carprofen (1/50 x body weight x 0.02) injected subcutaneously 60 min. before the surgery and kept anaesthetised with 2-4% isoflurane in a stereotactic frame to stabilized head. The cornea was incised near the junction with the sclera with a 25-G needle. Then, pre-loaded blunt cannula with islets was inserted into the ACE and islets were delivered inside. Eye ointment collyre containing erythromycin 0.5% was administered post-surgery.

### PA-mCherry adenovirus infection

Prior to transplantation into the ACE of recipients, isolated islets (from Ins1Cre:GCaMP6^f/f^ C57BL/6J donors, <12 weeks old) were infected with adenovirally-encoded (AV) PA-mCherry as previously described ^22^ for 24 h. This approach provided preferential infection of superficial beta cells (1-2 cells deep).

### *In vivo* Ca^2+^ imaging in the ACE

Ca^2+^ imaging was performed essentially as described ^21,29^.Imaging sessions were performed 21 weeks post-implantation under anesthesia with isoflurane (2-4%). All imaging experiments were conducted using a Zeiss LMS780 inverted confocal microscope equipped with 20x water dipping lenses (1.0 NA). The signal from GCaMP6f (ex. 488 nm, em. 525±25 nm) and PA-mCherry (ex. 541 nm, em. 570±25 nm) was recorded in time-series experiments for up to 30 min. Images were acquired for 2D at 3Hz (3 frames/s) and for 3D at 0.5Hz. To recover the same set of cells, PA-mCherry was utilized to provide “landmarks” to reorient the islet plane to the same as that of previous sessions. The focus was manually adjusted to correct for movement.

### Image analysis

Using FIJI (Image J), images in the time series were individually time-stamped to maintain their absolute time information before excluding frames where resolution was poor, or blurred by movement. Image series were then cropped and aligned using the plugin “Descriptor-based series registration (2d/3d + t)” (https://imagej.net/Descriptor-based_registration (2d/3d))38, applying the model “Rigid (2d)”, with “3-dimensional quadratic fit” ^32^. ROIs were created using GCaMP fluorescence and the negative shadow of nuclei. For each ROI-Cell, integrated fluorescence intensity and XY(Z) co-ordinates were compiled and processed for connectivity analysis. For longitudinal analysis, ROIs-Cells were manually matched across the videos.

### Connectivity analysis

Leader and hubs were identified as previously described ^22,30^. In each islet, the top 10% of cells with the shortest average time of activation per wave were considered as leaders. For hub cells, a threshold in which at least 80 % of connections presented connectivity of >0.8 were identified as highly-connected “hubs” ^30^.

### Statistical analysis

Statistical significance between two conditions was assessed using a paired or unpaired Student’s t-test. Interactions between multiple conditions were determined using one- or two-way ANOVA (with Tukey’s or Bonferroni posthoc tests). Analyses were performed using Graph Pad Prism (GraphPad Software version 8.0) and MATLAB (Mathworks) and significant p-values are described in each relevant section. Values are plotted as mean ± SEM, unless otherwise stated.

## RESULTS

### Leader cells are stable in vitro

We have recently reported that leader beta cells display a discrete transcriptome *versus* non-leaders and likely represent a stable population within the islet ^22^. We first explored Ca^2+^ dynamics and beta cell-beta cell connectivity *in vitro*, using islets in which the genetically-encoded Ca^2+^ sensor GCaMP was expressed selectively in these cells ^21^. This enabled us to identify leader cells from which Ca^2+^ waves emanated (SFig. 1). To confirm the stability of these cells *in vitro*, we used the cryptic fluorescent protein, PA-mCherry ^22^. Islets were transduced using adenovirus (*AV-CAG:PA-mCherry*) and 24h later, confocal imaging was performed (6Hz) (SFig. 1A-C, video 1). Temporally identified leader cells were “photo-painted” using UV light (SFig. 1F-G, video 2). Re-imaged 24h later, in 80% (8/10) of islets (*n*=4 mice), the labelled cell was identified as the leader cell again (SFig. 1E, F, video 3). However, we noted that during the second imaging session, the average time of response was shortened compared to the first, likely reflecting loss of functional performance related to collapse of the vasculature, cellular rearrangements, islet necrosis or other changes (SFig. 1, G-I).

### Ca^2+^ waves emanate from leaders and are transmitted by hubs *in vivo*

Given the challenges of studying the stability of individual cells *in vitro* (SFig. 1), we sought to establish a means of exploring beta cell functional stability in the living animal. To this end, we used wild-type animals as graft recipients, and Ins1Cre:GCaMP6f^fl/fl^ mice as islet donors. After engraftment into the ACE, islets become revascularized and innervated within 3-4 weeks ^33^. Using fast confocal imaging (3Hz), we recorded the Ca^2+^ dynamics of individual islets with cellular resolution while the animals were under anesthesia (see Methods). At this imaging speed, we were able to record the sequential activation of individual cells across a single 2D plane of the islet (Fig. 1A, video 4). In order to identify individual cells unambiguously, a projection of the time series was generated across the whole acquisition period (Fig. 1B). In line with earlier studies ^21,29^, spatially organised Ca^2+^ oscillations were remarkably stable over time during individual imaging sessions (typically of 12 min. duration; Fig. 1C) and in the face of steady blood glucose levels (12-16 mM; not shown). Ca^2+^ waves emanated from defined leader cells, present in discreet locations (Fig. 1A, arrows) and usually traversed the 2D imaging plane. Leader cells were formally defined as those in which the lowest average time of response (i.e. the time taken to increase fluorescence to 20% above the baseline for each Ca^2+^ wave; Fig. 1D). Deployment of previously described approaches ^21,30,34^ revealed network behaviours in which a substantial proportion of cells were highly connected (with a connectivity >0.8, in >70% of connections; Fig. 1E, “hubs”).

**Figure 1.**
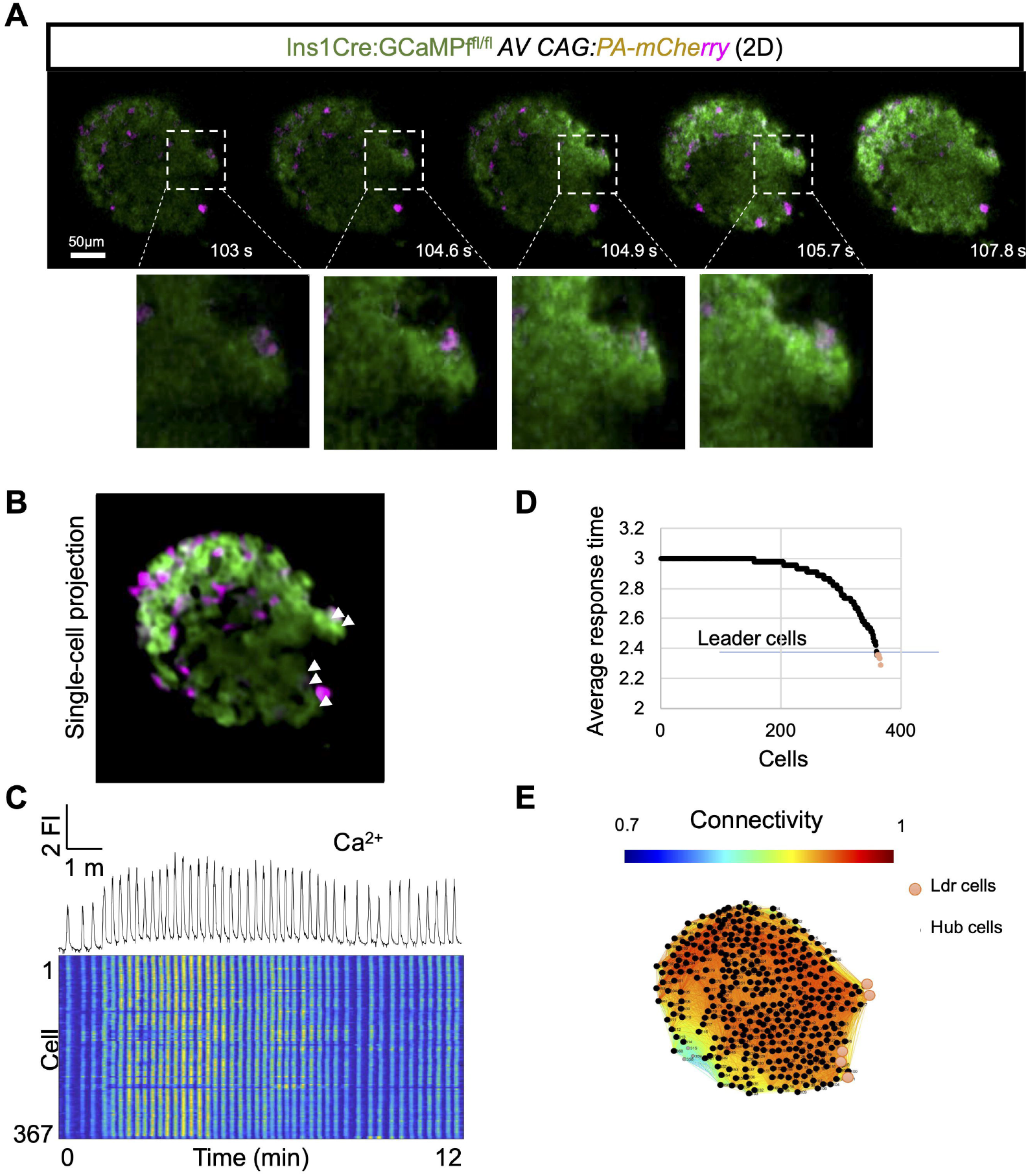
*In vivo* Ca^2+^ dynamics from engrafted islet into ACE. **A)** Snapshots from time series of confocal calcium imaging of pancreatic islet during a calcium wave at the indicated time points (in sec.) *in vivo*. The white box shows a group of “leader” cells. Scale bar = 50 µm. **B)** Time series projection used to identify individual cells from the imaged islet shown in (A). **C)** Fluorescent trace of the GCaMP6 signal from the islet shown in (B). The raster plot corresponds to the signal from individual cells. **D)** Average time of response for individual cells. The fastest top 10% are labelled as leader cells. **E)** Connectivity map and matrix of the islet shown in (A-B) indicating hub (black-dots) and leader (red-dots).

### Leader are persistent, but hub function is undermined by metabolic stress

We next explored the stability of both leaders and hubs *in vivo*. These parameters were examined both in wild type C57BL6 mice on a regular chow diet and in two models of type 2 diabetes: (i) metabolic stress induced by high fat-high sucrose (HFHS) feeding and the resultant obesity, and (ii) in animals in which one allele of the endogenous glucokinase (*Gck*) gene was disrupted by the insertion of cDNA encoding the fluorescent protein mCardinal, leading to alternative splicing (ca. 50% residual glucokinase activity; C57BL6/J-*Gck*^tm1(mCard)/*Rutt*^; *Gck*^*KI/+*^). As described in Fig. 2A, isolated islets from Ins1Cre:GCaMP6f^fl/fl^ mice infected with AV-CAG:PA-mCherry were engrafted into animals from three different groups: 1) wild type (WT) on regular chow diet, 2) WT on HFHS diet and 3) *Gck*^*KI/+*^ animals on chow diet. Each of thee two latter models displays mild hyperglycemia ^30,35^. 21 weeks post-engraftment, Ca^2+^ dynamics were recorded during anaesthesia at 0, 1 and 7 days (0d, 1d, 7d; Fig. 2B). We took advantage of the fact that while cells freshly infected with PA-mCherry were non-fluorescent for >24 h, after engraftment a random subset of cells spontaneously became fluorescent *in vivo* during the 21 week period before imaging. The fluorescent cells thus provided convenient “landmarks”, enabling identification of the same imaging plane, and thus the same set of cells, during sequential imaging sessions (Fig. 2Ci, Cii, Ciii, video 5-7).

**Figure 2.**
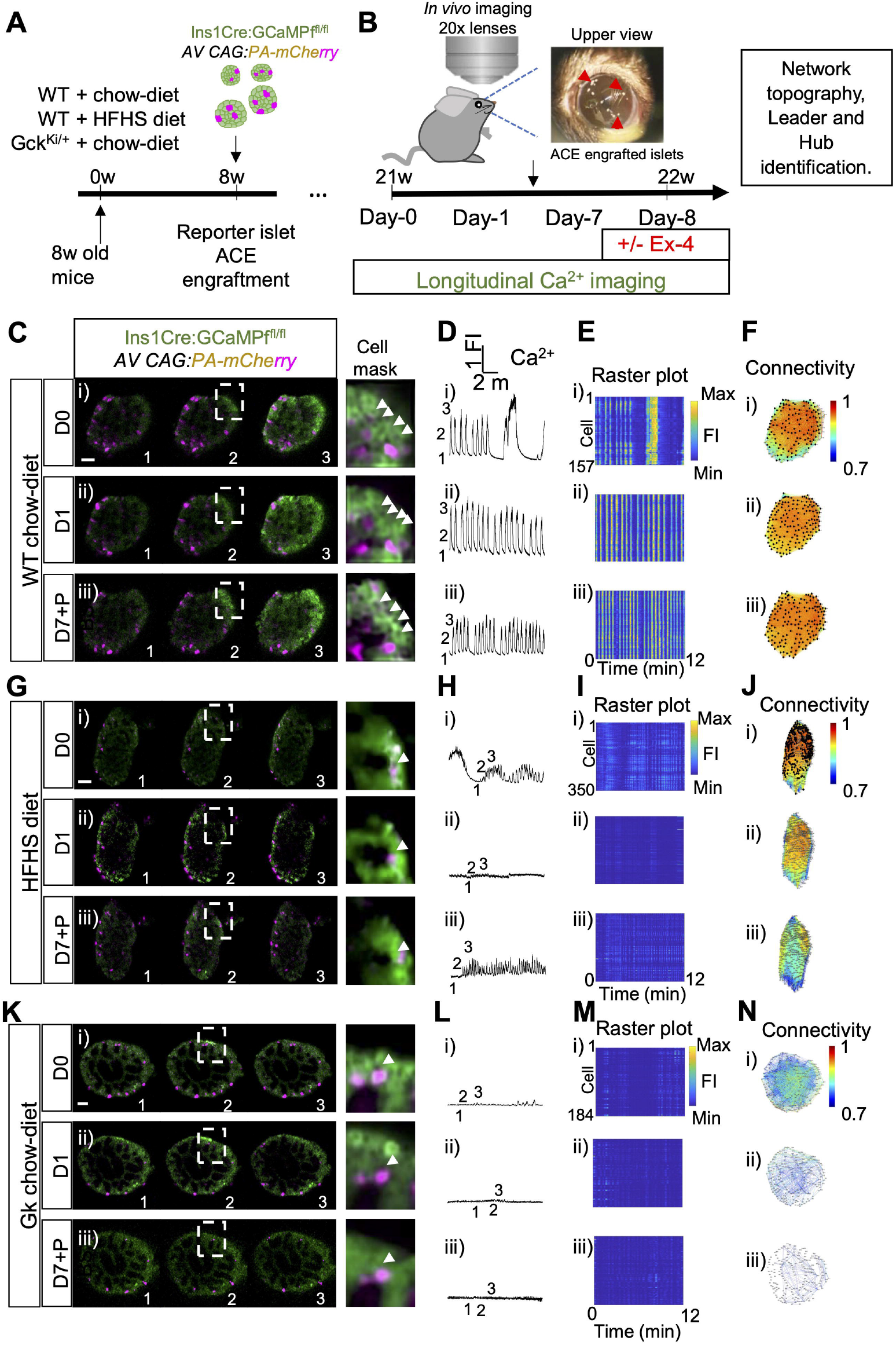
Leaders are stable in control, diet- and genetically-induced metabolic stress. **A)** Timeline of procedures. Animals were divided into three groups: WT + chow-diet, WT + HFHS-diet and Gck^KI/+^ + chow diet. After 8 weeks, isolated islets from WT Ins1Cre:GCaMP6^f/f^ were infected with AV CAG:PA-mCherry and engrafted into the anterior chamber of the mouse eye of each group. In these islets the genetically encoded Ca^2+^ indicator GCaMP6 (green) is specifically expressed in pancreatic beta cells and the PA-mCherry spontaneously activated in these engraftments served as landmarks. **B)** Ca^2+^ dynamics were recorded after 21 weeks under the indicated diets, in islets at day 0, 1 and 7 days. On day 8, an IP of vehicle or Exendin-4 was administered and Ca^2+^ dynamics recorded. After longitudinal recordings, “leaders” and “hubs” beta cells were identified (see Methods) and their islet network topographies were plotted. **C, G, K)** Snapshots from time series of confocal Ca^2+^ imaging of pancreatic islets at the indicated time points (in sec.) *in vivo*. The same islet was re-imaged at day 0, day 1 and day 7. The white arrowheads points to “leader” cells. Scale bar = 25 µm. **D, H, L)** Fluorescent traces of the GCaMP6 signal from the islets shown in (C, G, K). **E, I, M)** Raster plot corresponding to the signal from individual cells from the islets shown in (C, G, K). **F, J, N)** Connectivity maps of the islets shown in (E-G). In islets from WT animals on chow diet enjoy of highly coordinated activity, with Ca^2+^ waves repeatedly emanating from the same set of leader cells at different days. However, in WT animals fed with HFHS and Gck^Ki/+^ lower connectivity was observed, characterized by incomplete and abortive Ca^2+^ waves (Scale bar 50μm).

We found that in wild-type animals on a regular chow diet, islets displayed steady and spatially-organized Ca^2+^ waves at d0 under conditions in which blood glucose levels ranged from 12-16 mm during the recordings. The Ca^2+^ waves emanated from leader cells, which were consistently located at the islet periphery (Fig. 2Ci, Cii, Ciii). The waves were highly coordinated, and beta cells were highly connected (see Methods and Fig. 2Fi). However, and in agreement with an earlier study ^29^, islets engrafted into HFHS diet animals, or into mice heterozygous for the hypomorphic *Gck* allele (Methods), displayed impaired Ca^2+^ dynamics. Although leader cells were still detected, Ca^2+^ waves failed to be transmitted towards the rest of the islet (Fig 2Gi, video 8; 2Ki, video 9). Correspondingly, islets showed low connectivity (Fig 2Hi-Ji, 2Li-2Ni) and fewer hub cells than islets engrafted into in WT animals on regular chow. Blood glucose levels during imaging sessions were similar between control mice and animals on HFHS diet (10-16 mM in each case), but tended to be higher and more variable for *Gck*^*KI/+*^ mice (12-20 mM).

Animals from each group were reimaged 24h (1d) and 7d after the first inspection, using the PA-mCherry signal as a landmark to recover the same set of cells (see above). At both time points, we found that in controls, the same set of leaders and hubs were retrieved (Fig. 2C-F), demonstrating the stability of these cells. However, in both HFHS and *Gck*^*KI/+*^ groups, abortive and incomplete Ca^2+^ waves were observed which did not propagate across the islet (HFHS, Fig 2G-J, video 10-11; *Gck*^*Ki*^*/*, Fig 2K-N, video 12-13). Fewer hub cells were thus identified compared to control conditions. Corresponding blood glucose levels after Ex-4 injection were again similar between control mice, and animals on HFHS diet (9-16 mM), but tended to be higher in *Gck*^*KI/+*^ mice (10-20 mM). Thus, acute and genotype or incretin mimetic-dependent changes in blood glucose level are unlikely to underlie the differences in Ca^2+^ dynamics, leader / hub cell numbers, and connectivity.

### Acute GLP1R agonism reconnects beta cells across the islet network

We next explored the effects of glucagon-like peptide-1 receptor (GLP-1R) agonism on islet Ca^2+^ dynamics and connectivity *in vivo*. To this end, we first recorded Ca^2+^ dynamics at day 8 after IP injection of vehicle (PBS). Acute GLP1R agonism was then achieved by intraperitoneal injection of exendin-4 (Ex-4, 1 nmol/kg). Subsequent imaging allowed data capture within 5 min. of the injections. In WT chow diet-fed animals, injection of Ex-4 did not significantly affect connectivity, nor the percentage of detected hubs (Fig. 3A-D, 3M, 3N, video 14). In contrast, in HFHS and *Gck*^*KI+*^ animals, acute IP injection of Ex-4 rescued defective islet Ca^2+^ dynamics (HFHS, Fig 2E-H, video 15; *Gck*^*KI+*^,Fig 2I-L, video 16). Notably, Ex-4 reengaged a hub cell population, restoring beta cell-beta cell connectivity and Ca^2+^ dynamics (Fig. 3M-3N) (Ctrl + PBS versus Ctrl+Ex-4 not significant, *n*=5, HFD+PBS vs HFD+Ext-4 p=0.011, *n*=4, *Gck*^*KI/+*^+PBS vs *Gck*^*KI/+*^+Ext-4, *p*=0.026, *n*=5; Fig. 3E-L, 3M, 3N).

**Figure 3.**
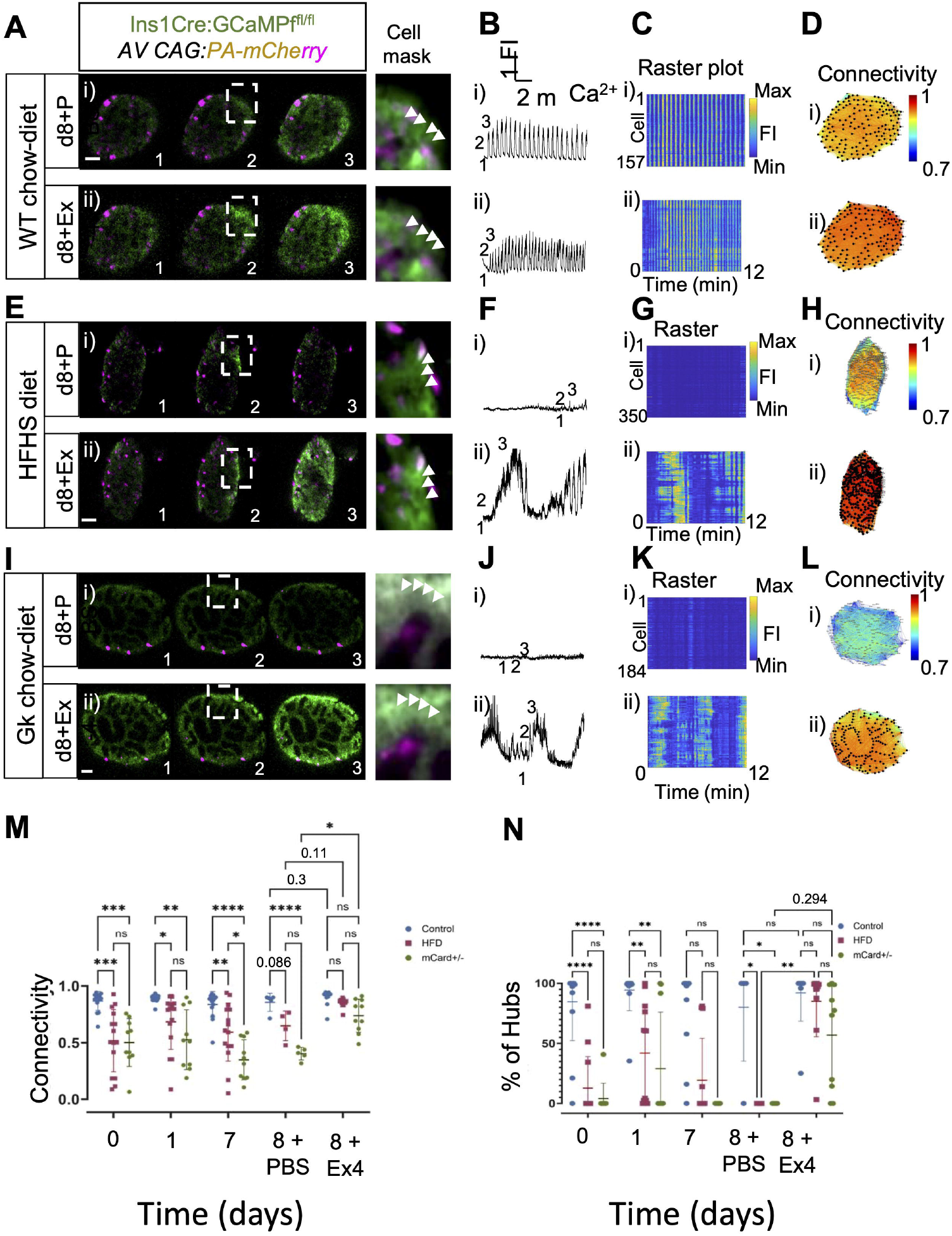
Connectivity is restored after acute Exendin-4 injection. **A, E, I)** Snapshots from timeseries of confocal calcium imaging of pancreatic islet showing Ca^2+^ waves at the indicated time points (in sec.) *in vivo*. The same islet was recorded first, upon injection of PBS, then upon injection of Exendin-4. The white box shows the area containing the “leader” cells (cell mask). Scale bar = 25 µm. **B, F, J)** Fluorescent traces of the GCaMP6 signal from the islets shown in (A, E, I) **C, G, K)** The Raster plot corresponds to the signal from individual cells from the islets shown in (A). **D, H, L)** Connectivity maps of the islets shown in (A, E, I). In control islets, there was no change in islet activity. However, in HFHS fed WT and Gck^Ki/+^ Exendin-4 administration restored islet Ca^2+^ dynamics, mainly by restoring “hub” function. **M)** Connectivity quantification from each group at day 0, 1, 7(+PBS), 8(+PBS) and 8(+Ex-4). **N)** Plot showing the average percentage of hubs from each group at day 0, 1, 7(+PBS), 8(+PBS) and 8(+Ex-4). Of note, upon Exendin-4 administration on day 8, hub cells reemerged (Scale bar 50μm).

### 3D recordings demonstrate that GLP1R agonism reconnects the entire islet

We next asked whether the above observations made across a single imaging plane in 2D could be extended across the islet in three dimensions. We therefore collected images every 2s, in imaging planes spaced 10-15 µm apart, firstly in the WT, regular chow diet condition (Fig. 4 A-F, video 17). As observed in 2D, we observed highly coordinated activity across the islet. Leaders and hubs were identified using the same approaches as in 2D. Importantly, no significant Ex-4-induced differences in connectivity or overall activity were observed across the imaged planes (Fig. 4Gi-Gii, video 18). However, small Ex4-induced changes in oscillation frequency or wave width were apparent (SFig. 3). In contrast, but consistent with 2D imaging (Fig. 3A,E), when islets were imaged in 3D, intraperitoneal injection of Ex-4 into HFHS diet animals led to the re-emergence of hub and (to a lesser extent) leader cells (Fig. 4 H-N, video 19).

**Figure 4.**
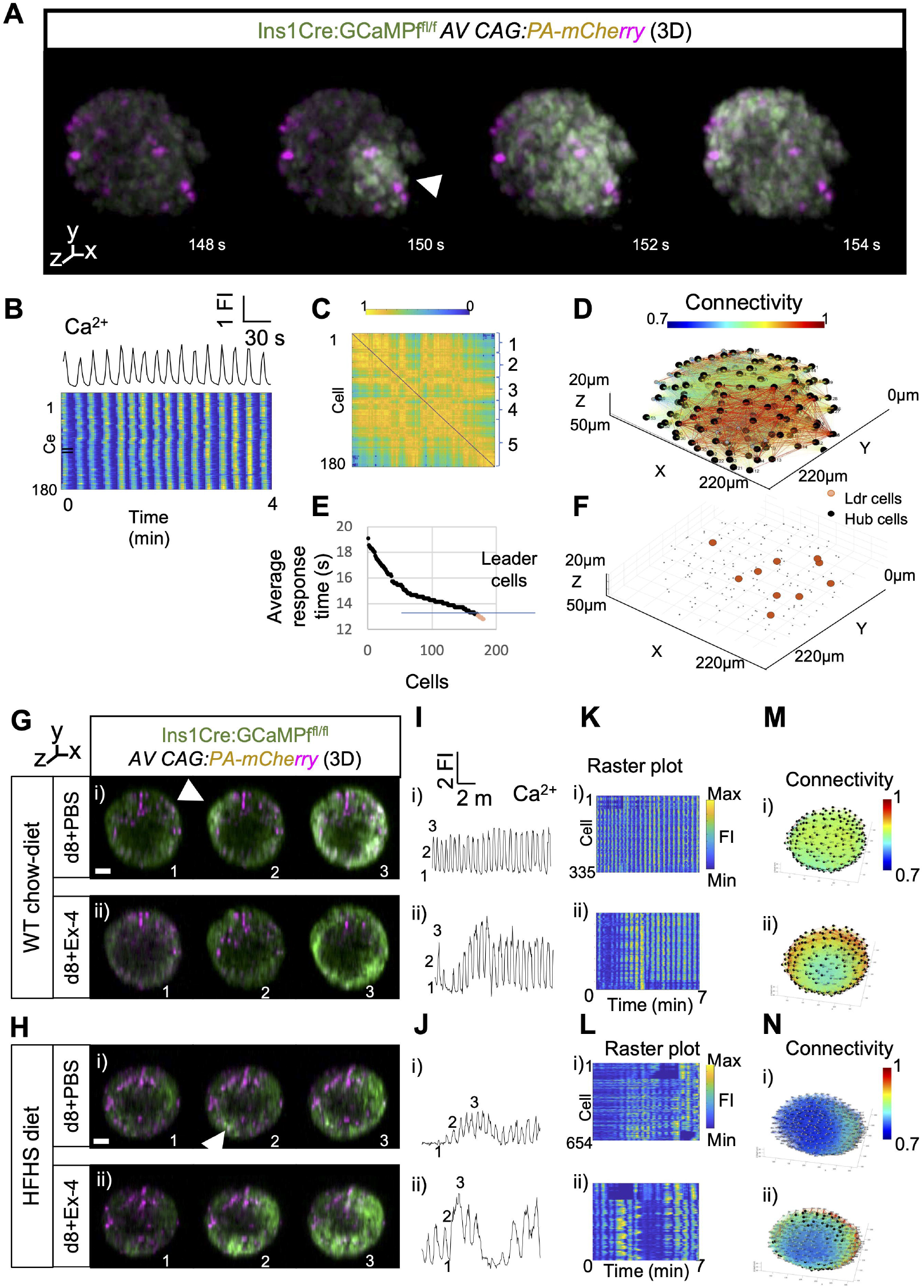
Connectivity is restored after acute Exendin-4 exposure: analysis in 3D. **A)** 3D projections from timeseries of confocal calcium imaging of pancreatic islet. Scale bar = 50 µm in XY and Z-depth. **B)** Fluorescent trace of the GCaMP6 signal from the islet shown in (A) The Raster plot corresponds to the signal from individual cells. **C)** Matrix showing the individual connectivity across the islet with the right braces indicating the different planes recorded. **D)** 3D connectivity topography showing the hub cells. **E)** Average time of response for individual cells. The fastest top 10% are labelled as leader cells. **F)** Cell map of the islet shown in (B), indicating the leaders (red-dots). **G-H)** 3D projections from timeseries of confocal Ca^2+^ imaging of pancreatic islet showing a Ca^2+^ waves at the indicated time points (in sec.) *in vivo*. The same islet was recorder first, upon injection of vehicle, then upon injection of Exendin-4. The white arrowheads point at the “leader” cells. Scale bar = 100 µm. **I-J)** Fluorescent traces of the GCaMP6 signal from the islets shown with vehicle or Exendin-4 IP. **K-L)** Raster plot corresponds to the signal from individual cells. **M-N)** Connectivity maps of the islets shown in (G-H); in control islets, there was an increase on 3D connectivity. In WT animals with HFHS diet, Exendin-4 administration restored islet Ca^2+^ calcium dynamics, and “hub” cells re-emerged (Scale bar 50μm).

## Discussion

The chief goals of the present study were: (1) to develop an approach to monitor the functional roles of *individual* beta cells within the islet syncytium *in vivo* over time, and (2) and to explore how the behaviours of these cells may change in models of type 2 diabetes and in response to a diabetes therapy (GLP-1R agonism). Our findings reveal that the apparent functional “identity” of cell populations is remarkably stable *in vivo* under normoglycemic conditions over time (one week), demonstrating that the existence of discrete populations at any given time point does not simply reflect stochastic events or ephemeral “substates” ^36^. Importantly, we show that functional compromise at high glucose levels conditions is rapidly rescued, independently of changes in glucose, by incretin receptor engagement. The intrinsic properties of these cell populations thus remain even when they have been subjected to chronic exposure to a hyperglycemic milieu.

We demonstrate firstly that Ca^2+^ waves emanate from leader cells that are stable for at least seven days. In contrast, hub cells display higher variability with ∼77% stable (*n*=8). Interestingly, we noted that whereas leaders were usually located towards the islet periphery, hubs were randomly distributed across the islet, whether explored in 2D or in 3D. Importantly, we did not note any greater preponderance of hubs in the proximity of blood vessels, whose locations were readily identified as dark areas in the islet images and movies.

Next, we reveal the effect of long-term metabolic stress examining islets engrafted into animals fed with high-fat-high-sucrose (HFHS) diet or into mice heterozygous for a hypomorphic *Gck* allele (*Gck*^*KI/+*^) which display mild hyperglycemia ^30^. In each of these disease models, leader cells were still clearly evident, whilst hub cell numbers were substantially lowered, with islets displaying impaired Ca^2+^ dynamics and lower connectivity (e.g. incomplete and abortive Ca^2+^ waves). In each setting, acute GLP1R agonism with Ex-4 restored islet Ca^2+^ dynamics within minutes in both HFHS and *Gck*^*KI+*^ animals, reengaging a “hub” population to restore beta cell-beta cell connectivity. We note that this rescue was associated with minimal changes acutely (usually a small *decrease)* in glucose concentration (from 12-16mM after PBS or 10-16mM Ex-4 injection), which are thus unlikely to explain the acute enhancement of beta cell function. Importantly, whereas 2D imaging allowed us to explore connectivity and “leadership” with high temporal resolution (3Hz), 3D imaging allowed islet-wide connectivity to be explored albeit at the cost of temporal and axial resolution (0.5Hz). Nevertheless, we deemed the latter strategy important since it provided both a demonstration of islet-wide reconnection in response to GLP1R agonisms, without substantial compromise of connectivity analysis which may occur at lower acquisition speeds.

Of note, the effects of incretin receptor agonism were more substantial than we ^34,37^ and others ^38^ have previously reported *in vitro* at least in islets isolated from islets from non-obese or hyperglycemic mice. Interestingly, whereas Roger et al ^38^ reported the elimination of GLP-1 responses after incubation of islets at elevated glucose concentrations, we observed that the effects of agonism at these receptors *in vivo* were modest in normoglycemic mice (SFig. 3), but dramatic when examined in two models of hyperglycemia (Fig. 3). Although requiring further exploration, our results therefore suggest that GLP1RA may act at sites outside of the islet, possibly through neuronal relays (e.g. from the hind brain, paraventricular nuclei, gut or other locations where GLP1R are enriched) ^39^, providing possible parasympathetic stimulation of the islet ^40^. Further studies using pharmacological and surgical approaches will be required to explore these possibilities.

### Limitations of the study

We have explored islet function after engraftment in the anterior eye chamber, allowing the preservation of beta cell function of days, and providing physiological exposure to neuronal and humoral factors. We note that whilst parasympathetic input into the eye – via the oculomotor nerve ^41^ – is likely to recapitulate, at least in large part, that seen in in the pancreas, we do not exclude subtle differences between the two, or the possibility that those innervating the eye ^42^ may arise from areas of the brain more enriched for GLP1R.

The present longitudinal study was limited to eight days. Whether the observed cellular stabilities persist over longer time periods is unclear, and future studies will be needed to explore this question. We note that changes in islet cellularity (hypertrophy and hyperplasia) in response to metabolic stress ^43^ may provide a challenge to such analyses. Moreover, the present studies were confined to male mice. Whether the observations can be extended to females will need to be explored.

The studies here used only a single GLP1RA, Exendin-4, a product of the Gila monster with a drastically extended half-life in the circulation compared with native GLP1R ^44^. However, use of Exendin-4 (also called exenatide) in the clinic is declining, with stabilised GLP1R agonists such as liraglutide and semaglutide more usually deployed to treat T2D ^45^. Dual and triple receptor agonists (interacting additionally with glucose-dependent insulinotropic peptide, GIP or glucagon receptors) are increasing deployed in patients to treat both obesity and diabetes ^46,47^, and their impact on Ca^2+^ dynamics, as described here, would be of interest.

### Conclusions and perspectives

We describe a “landmark labelling” approach through which individual pancreatic beta cell function and connectivity can be monitored with cellular resolution *in vivo*. We use this technique to show how these parameters are affected reversibly in disease states and by treatment with an anti-hyperglycemic agent. Critically, we demonstrate that individual beta cells with highly specialised functions (leaders, hubs) retain these roles over many days in normoglycemic animals. The loss of hub, but not leader, function is the chief feature of two models type 2 diabetes. Remarkably, we show that the former role re-emerges within minutes of treatment with an incretin mimetic hormone.

Our strategy, which combines cellular labelling, functional imaging of Ca^2+^ dynamics through the front of the eye, and prospective imaging over many days in the same animal, may prove useful to explore the functional evolution and stability of individual cells in other disease states.

## CRediT author statement

**LD**, conceptualisation, methodology, experimentation, analysis; **SS**, investigation, analysis; **LLN, APG, SL**, experimentation; **AP**, methodology; **GR**, conceptualisation, writing, funding, supervision.

---

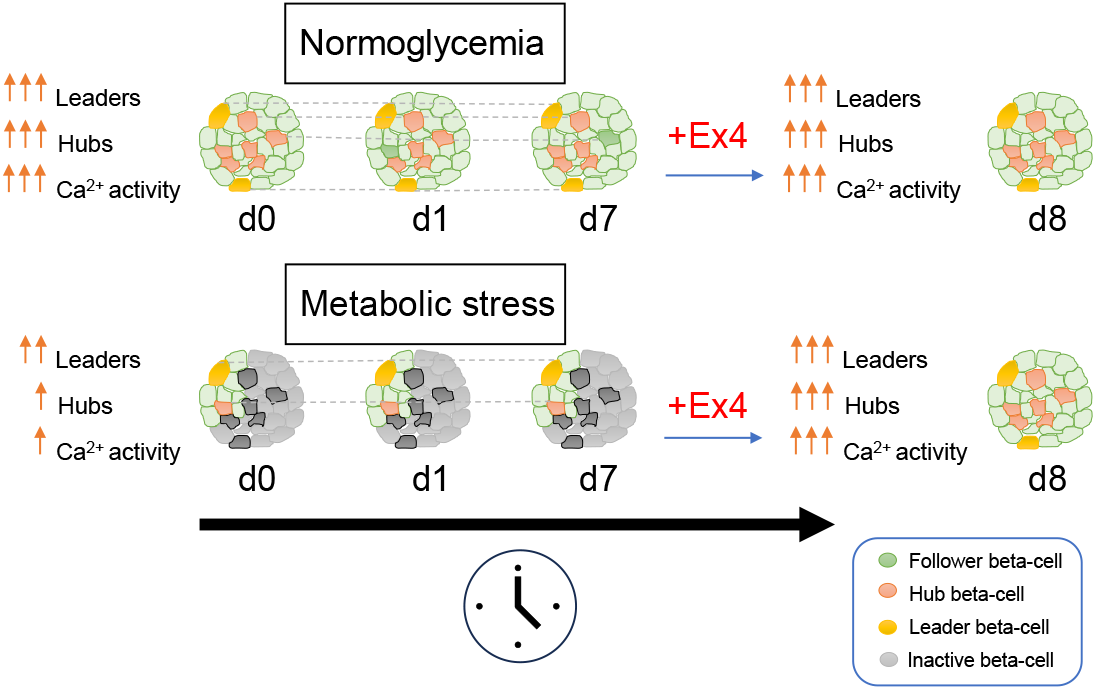

## References

1. Eisenbarth, G.S. Banting Lecture 2009: An unfinished journey: molecular pathogenesis to prevention of type 1A diabetes. Diabetes 59, 759–74 (2010).

2. Suzuki, K. et al. Genetic drivers of heterogeneity in type 2 diabetes pathophysiology. Nature 627, 347–357 (2024).

3. Cohrs, C.M. et al. Dysfunction of Persisting beta Cells Is a Key Feature of Early Type 2 Diabetes Pathogenesis. Cell Rep. 31, 107469 (2020).

4. Müller Wa Fau-Faloona, G.R., Faloona Gr Fau-Aguilar-Parada, E., Aguilar-Parada E Fau-Unger, R.H. & Unger, R.H. Abnormal alpha-cell function in diabetes. Response to carbohydrate and protein ingestion.

5. O’Rahilly, S., Turner, R.C. & Matthews, D.R. Impaired pulsatile secretion of insulin in relatives of patients with non-insulin-dependent diabetes. N.Engl.J.Med. 318, 1225–1230 (1988).

6. Rutter, G.A., Sidarala, V., Kaufman, B.A. & Soleimanpour, S.A. Mitochondrial metabolism and dynamics in pancreatic beta cell glucose sensing. Biochem J 480, 773–789 (2023).

7. Rorsman, P. & Ashcroft, F.M. Pancreatic beta-Cell Electrical Activity and Insulin Secretion: Of Mice and Men. Physiol Rev. 98, 117–214 (2018).

8. Merrins, M.J., Corkey, B.E., Kibbey, R.G. & Prentki, M. Metabolic cycles and signals for insulin secretion. Cell Metab 34, 947–968 (2022).

9. Rutter, G.A. & Sweet, I.R. Glucose Regulation of β-Cell KATP Channels: Is a New Model Needed? Diabetes 73, 849–855 (2024).

10. Leech, C.A. et al. Molecular physiology of glucagon-like peptide-1 insulin secretagogue action in pancreatic beta cells. Prog.Biophys.Mol.Biol. 107, 236–247 (2011).

11. Nadal, A., Quesada, I. & Soria, B. Homologous and heterologous asynchronicity between identified alpha-, beta- and delta-cells within intact islets of Langerhans in the mouse. J Physiol 517, 85–93 (1999).

12. Salomon, D. & Meda, P. Heterogeneity and contact-dependent regulation of hormone secretion by individual B-cells. Exp.Cell Res. 162, 507–520 (1986).

13. Kiekens, R. et al. Differences in glucose recognition by individual rat pancreatic B cells are associated with intercellular differences in glucose-induced biosynthetic activity. J.Clin.Invest. 89, 117–125 (1992).

14. Van Schravendijk, C.F., Kiekens, R. & Pipeleers, D.G. Pancreatic beta cell heterogeneity in glucose-induced insulin secretion. J.Biol.Chem. 267, 21344–21348 (1992).

15. Dorrell, C. et al. Human islets contain four distinct subtypes of beta cells. Nat.Commun. 7:11756. doi: 10.1038/ncomms11756., 11756 (2016).

16. Wang, Y.J. et al. Single-Cell Transcriptomics of the Human Endocrine Pancreas. Diabetes. 65, 3028–3038 (2016).

17. Segerstolpe, A. et al. Single-Cell Transcriptome Profiling of Human Pancreatic Islets in Health and Type 2 Diabetes. Cell Metab. 24 593–607 (2016).

18. Yu, V. et al. Differential CpG methylation at Nnat in the early establishment of beta cell heterogeneity. Diabetologia 67, 1079–1094 (2024).

19. Singh, S.P. et al. Different developmental histories of beta-cells generate functional and proliferative heterogeneity during islet growth. Nat.Commun. 8, 664–00461 (2017).

20. Johnston, N.R. et al. Beta cell hubs dictate pancreatic islet responses to glucose. Cell Metabolism 24, 389–401 (2016).

21. Salem, V. et al. Leader beta cells coordinate Ca2+ dynamics across pancreatic islets in vivo. Nat Metab. 1, 615–629 (2019).

22. Chabosseau, P. et al. Molecular phenotyping of single pancreatic islet leader beta cells by “Flash-Seq”. Life Sci 316, 121436 (2023).

23. Delgadillo-Silva, L.F. et al. Optogenetic β cell interrogation in vivo reveals a functional hierarchy directing the Ca(2+) response to glucose supported by vitamin B6. Sci Adv 10, eado4513 (2024).

24. Gosak, M. et al. Critical and Supercritical Spatiotemporal Calcium Dynamics in Beta Cells. Front Physiol. 8, 1106 (2017).

25. Westacott, M.J., Ludin, N.W.F. & Benninger, R.K.P. Spatially Organized beta-Cell Subpopulations Control Electrical Dynamics across Islets of Langerhans. Biophys.J. 113, 1093–1108 (2017).

26. Kravets, V. et al. Functional architecture of pancreatic islets identifies a population of first responder cells that drive the first-phase calcium response. PLoS Biol 20, e3001761 (2022).

27. Rutter, G.A., Gresch, A., Delgadillo Silva, L. & Benninger, R.K.P. Exploring pancreatic beta-cell subgroups and their connectivity. Nat Metab 6, 2039–2053 (2024).

28. Szabat, M. et al. Kinetics and genomic profiling of adult human and mouse β-cell maturation.

29. Akalestou, E. et al. Intravital imaging of islet Ca(2+) dynamics reveals enhanced ×ý cell connectivity after bariatric surgery in mice. Nat Commun. 12, 5165 (2021).

30. Salazar, S. et al. Sex-dependent additive effects of dorzagliatin and incretin on insulin secretion in a novel mouse model of GCK-MODY. bioRxiv (2024).

31. Ravier, M.A. & Rutter, G.A. Isolation and culture of mouse pancreatic islets for ex vivo imaging studies with trappable or recombinant fluorescent probes. Methods Mol.Biol. 633, 171–184 (2010).

32. Preibisch, S., Saalfeld, S., Schindelin, J. & Tomancak, P. Software for bead-based registration of selective plane illumination microscopy data. Nat.Methods. 7, 418–419 (2010).

33. Speier, S. et al. Noninvasive in vivo imaging of pancreatic islet cell biology. Nat.Med. 14, 574–578 (2008).

34. Hodson, D.J. et al. Lipotoxicity disrupts incretin-regulated human beta cell connectivity. J.Clin.Invest. 123, 4182–4194 (2013).

35. Muniangi-Muhitu, H. et al. Covid-19 and Diabetes: A Complex Bidirectional Relationship. Front Endocrinol.(Lausanne) 11, 582936 (2020).

36. Chu, C.M.J. et al. Dynamic Ins2 Gene Activity Defines β-Cell Maturity States. Diabetes 71, 2612–2631 (2022).

37. Cane, M.C., Parrington, J., Rorsman, P., Galione, A. & Rutter, G.A. The two pore channel TPC2 is dispensable in pancreatic beta-cells for normal Ca(2)(+) dynamics and insulin secretion. Cell Calcium. 59, 32–40 (2016).

38. Roger, B. et al. Adenylyl cyclase 8 is central to glucagon-like peptide 1 signalling and effects of chronically elevated glucose in rat and human pancreatic beta cells. Diabetologia. 54, 390–402 (2011).

39. Kabahizi, A. et al. Glucagon-like peptide-1 (GLP-1) signalling in the brain: From neural circuits and metabolism to therapeutics. Br J Pharmacol 179, 600–624 (2022).

40. Rodriguez-Diaz, R. et al. Noninvasive in vivo model demonstrating the effects of autonomic innervation on pancreatic islet function. Proc.Natl.Acad.Sci.U.S.A. 109, 21456–21461 (2012).

41. Jackson, P.C. Innervation of the iris by individual parasympathetic axons in the adult mouse. J Physiol 378, 485–95 (1986).

42. Landis, S.C., Jackson, P.C., Fredieu, J.R. & Thibault, J. Catecholaminergic properties of cholinergic neurons and synapses in adult rat ciliary ganglion. J Neurosci 7, 3574–87 (1987).

43. Chen, C. et al. Alterations in beta-Cell Calcium Dynamics and Efficacy Outweigh Islet Mass Adaptation in Compensation of Insulin Resistance and Prediabetes Onset. Diabetes. 65, 2676–2685 (2016).

44. Doyle, M.E. & Egan, J.M. Glucagon-like peptide-1. Recent Prog Horm Res 56, 377–99 (2001).

45. Nauck, M.A. & Müller, T.D. Incretin hormones and type 2 diabetes. Diabetologia 66, 1780–1795 (2023).

46. Madsbad, S. & Holst, J.J. The promise of glucagon-like peptide 1 receptor agonists (GLP-1RA) for the treatment of obesity: a look at phase 2 and 3 pipelines. Expert Opin Investig Drugs 34, 197–215 (2025).

47. Manchanda, Y. et al. Binding Kinetics, Bias, Receptor Internalization and Effects on Insulin Secretion in vitro and in vivo of a Novel GLP-1R/GIPR Dual Agonist, HISHS-2001. bioRxiv (2025).

