## Supplemental Figures for "Exploration of individual beta cell function over time *in vivo:* effects of hyperglycemia and glucagon-like peptide-1 receptor (GLP1R) agonism"

### Supplementary figures

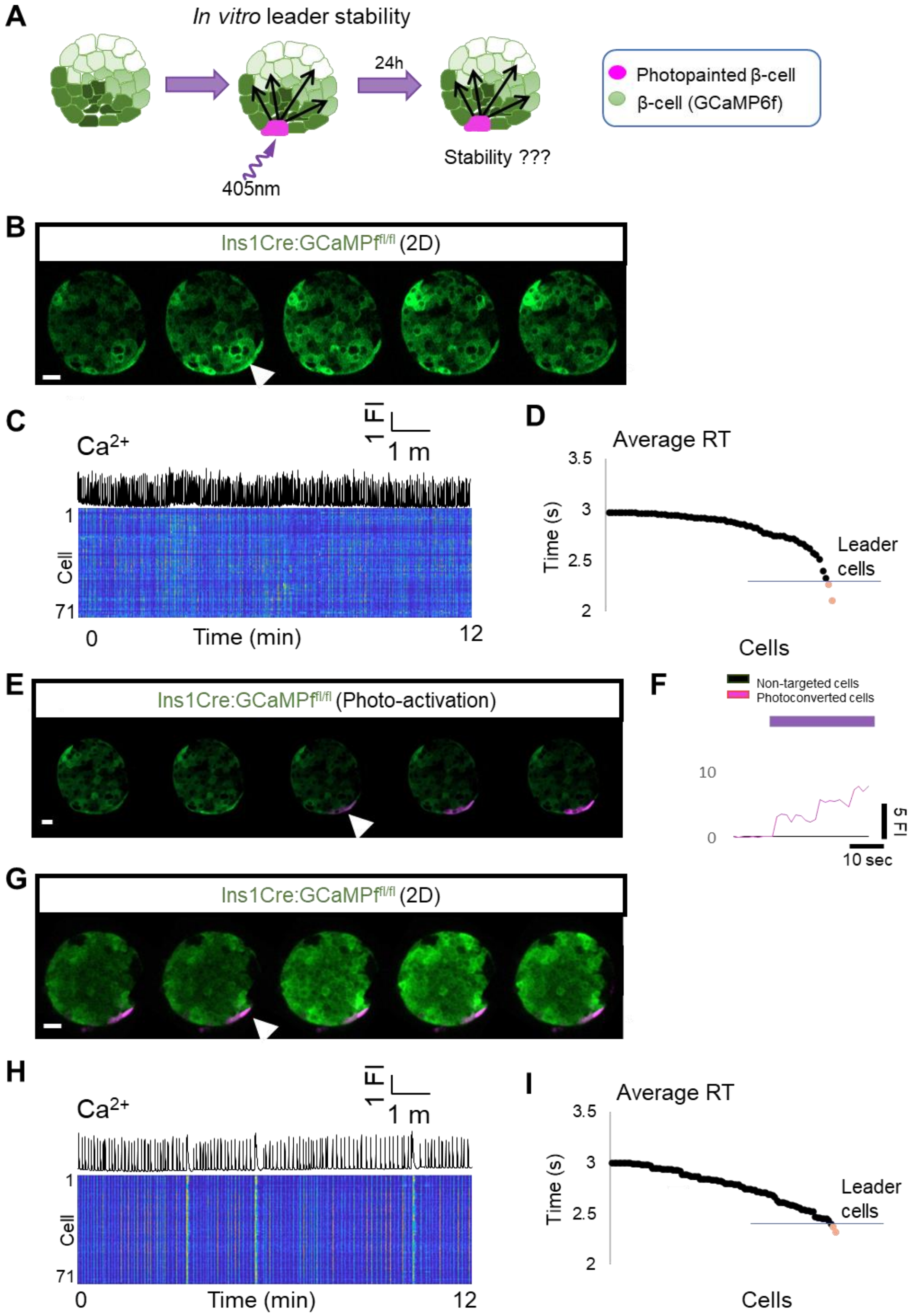


Supp. Fig 1

**Supplementary Figure 1. Leader cells are stable *in vitro*. A)** Cartoon showing the principle of short-lineage tracing by photopainting: in isolated islets infected with AV CAG:PA-mCherry, the leader cells are identified and photolabelled *in situ*. The islets are re-imaged 24hrs later to investigate the stability of leader cells. **B)** Snapshots from confocal images of an islet. **C)** Calcium traces from the islet shown in (A). The rasterplot shows the activity of individual cells. **D)** connectivity matrix and connectivity map showing the leader cells. **E)** Average time of response for individual cells of the islet shown in (A). The leader cells display the fastest time of response. **F)** Snapshots from the islet shown in (A). During the imaging, the leader cell is photopainted by FRAP (405nm). **G)** fluorescent traces of the ratio between the mCherry/GCaMP signal upon PA-mCherry activation. The photolabelled cell increases the mCherry signal while non-targeted cells remain unlabelled (black line). **H)** Snapshots from confocal images of an islet show in (A) and photpainted on (H). **J)** Connectivity matrix and connectivity map showing the leader cells. **I)** The photopainted leaders remained stable after 24h (Scale bar = 25µm).

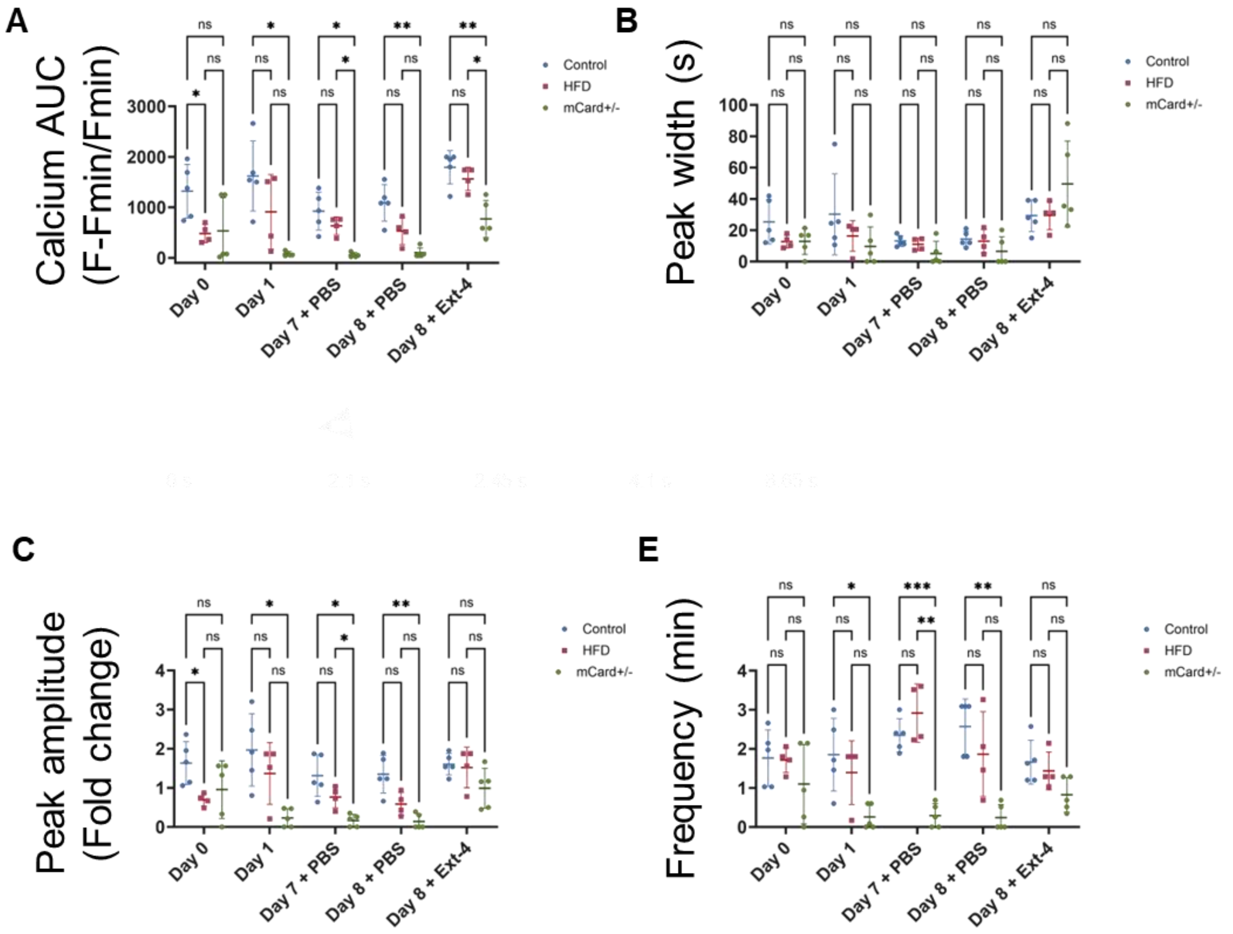


**Supplementary Figure 2. Exendin-4 improves islet function. A)** Quantifications of calcium AUC from 1) WT + chow-diet, 2) WT + HFHS-diet and 3) $Gck^{KI/+}$ + chow diet at day 0, 1, 7(+PBS), 8(+PBS) and 8(+Ex-4). **B)** Average peak width, **C)** peak amplitude and **E)** frequency of the imaged islets (n=4 for each group). (2-way paired ANOVA, Tukey's correction *P ≤ .05, **P ≤.01, ***P ≤ .001, ****P ≤ .0001, ns, not significant; n=3, WT + Chow diet n=4, WT + HFHS diet n=4, $Gck^{KI/+}$ + chow diet n=4. Data are mean ± SD.)

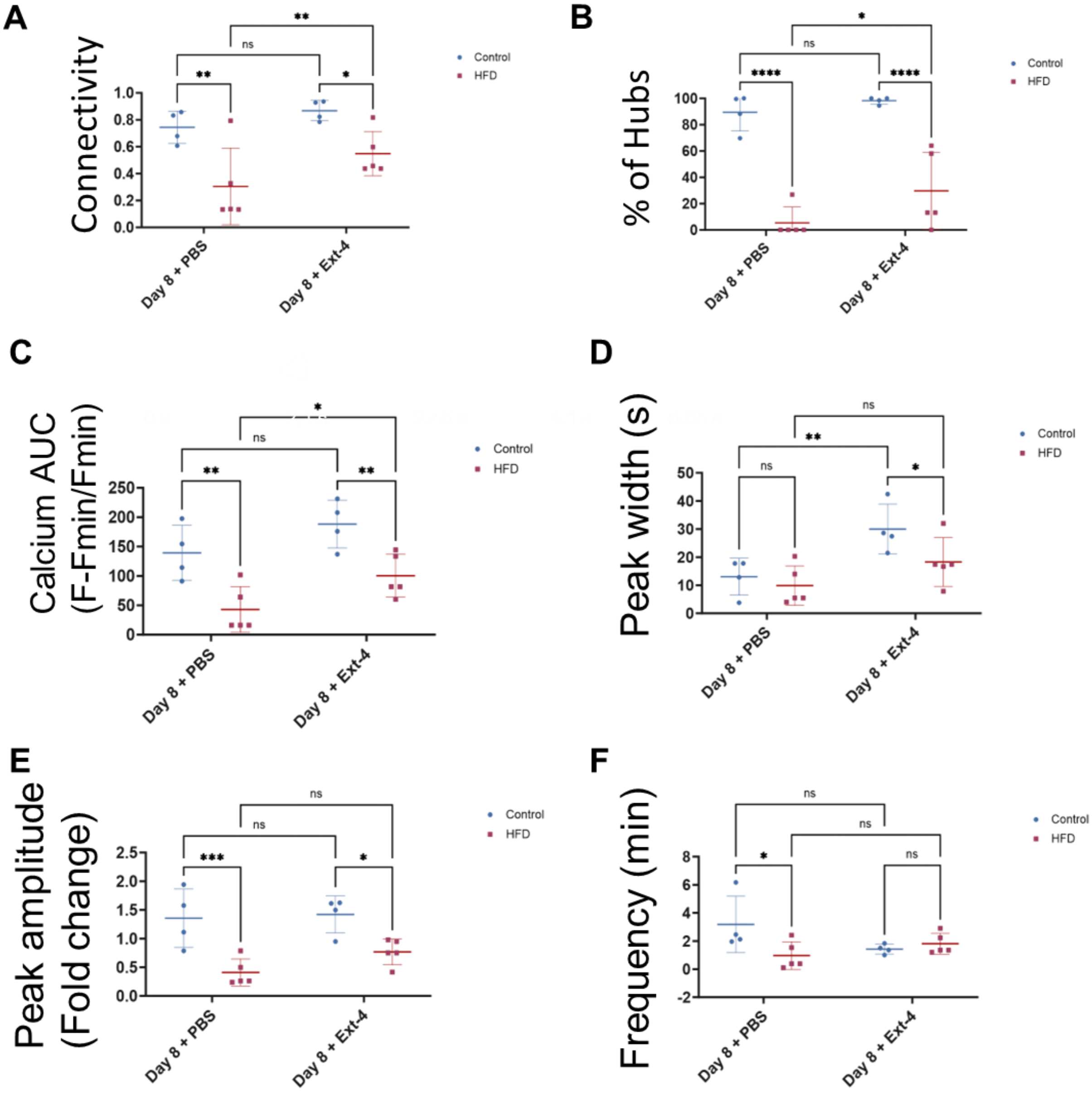


**Supplementary Figure 3. Exendin-4 improves islet-wide function. A)** Connectivity quantification 1) WT + chow-diet, 2) WT + HFHS-diet groups at day 8(+PBS) VS 8(+Ex-4) from 3D recordings. **B)** Percentage of highly connected cells, i.e. hubs, detected across the islet. **C)** Quantifications of calcium AUC. **D)** Average peak width, **E)** peak amplitude and **F)** frequency of the imaged islets (n=4 for each group). (2-way paired ANOVA, Tukey's correction *P ≤ .05, **P ≤.01, ***P ≤ .001, ****P ≤ .0001, ns, not significant; n=3, WT + Chow diet n=4, WT + HFHS diet n=5. Data are mean ± SD.)

**Supplementary video 1. Leader beta cells can be identified in vitro.** *In vitro* calcium imaging from transgenic C57BL/6J Ins1Cre:GCaMP6f$^{fl/fl}$ isolated islet. The video was recorded at 6Hz with constant 11mM glucose. The cells were the calcium wave initiates present the lowest average time of activation i.e the time to present an increase of >20% over the baseline over the calcium waves.

**Supplementary video 2. Leader beta cells photolabeling.** Photoactivation of the PA-mCherry using UV-light (λ = 405) (*n*=10).

**Supplementary video 3. Leader beta cells are stable in vitro.** *In-vitro* re-imaging after 24hrs of leader cell photolabeling. The photolabelled leader cell had the lowest average time of activation in 8/10 islets.

**Supplementary video 4. Intravital imaging of islets engrafted into the anterior chamber of the mouse eye.** *In vivo* calcium imaging of a transgenic C57BL/6J Ins1Cre:GCaMP6f$^{fl/fl}$ islet engrafted into a WT mouse. The islet Islets were transduced using adenovirus (*AV-CAG:PA-mCherry*). The video was recorded at 3Hz with blood glucose range from 12-16mM glucose. Leader cells can be identified *in vivo* i.e the cells that show the fastest response time.

**Supplementary video 5. Longitudinal Intravital imaging of islets engrafted into the anterior chamber of the mouse eye.** *In vivo* calcium imaging at day 0 after 21w post engraftment (see Methods) of a WT animal exposed to regular chow diet.

**Supplementary video 6. Longitudinal Intravital imaging of islets engrafted into the anterior chamber of the mouse eye.** *In vivo* calcium imaging after 24h (second imaging session) of the same islet.

**Supplementary video 7. Longitudinal Intravital imaging of islets engrafted into the anterior chamber of the mouse eye.** *In vivo* calcium imaging after 7 days (third imaging session) of the same islet engrafted into a WT animal exposed to regular chow diet.

**Supplementary video 8. Longitudinal Intravital imaging of islets engrafted into the anterior chamber of the mouse eye fed with HFHS diet.** *In vivo* calcium imaging at day 0 after 21w post engraftment (see Methods) of a WT animal exposed to high fat high sugar (HFHS) diet. The islet displayed incomplete and abortive calcium waves.

**Supplementary video 9. Longitudinal Intravital imaging of islets engrafted into the anterior chamber of the mouse eye of *Gck*$^{Ki/+}$ animals.** *In vivo* calcium imaging at day 0 after 21w post engraftment (see Methods) of a Gck$^{Ki/+}$ animal exposed to regular chow diet.

**Supplementary video 10. Longitudinal Intravital imaging of islets engrafted into the anterior chamber of the mouse eye, fed with HFHS diet.** *In vivo* calcium imaging after 24h (second imaging session) of the same islet (HFHS diet).

**Supplementary video 11. Longitudinal Intravital imaging of islets engrafted into the anterior chamber of the mouse eye, fed with HFHS diet.** *In vivo* calcium imaging after 7 days (third imaging session) of the same islet (HFHS diet).

**Supplementary video 12. Longitudinal Intravital imaging of islets engrafted into the anterior chamber of the mouse eye of *Gck*$^{Ki/+}$ animals.** *In vivo* calcium imaging after 24h (second imaging session) of the same islet engrafted into Gck$^{Ki/+}$ animal.

**Supplementary video 13. Longitudinal Intravital imaging of islets engrafted into the anterior chamber of the mouse eye of *Gck*$^{Ki/+}$ animals.** *In vivo* calcium imaging after 7 days (third imaging session) of the same islet engrafted into Gck$^{Ki/+}$ animal.

**Supplementary video 14. Longitudinal Intravital imaging of islets engrafted into the anterior chamber of the mouse eye.** *In vivo* calcium imaging at day 8 after vehicle (PBS) injection followed by a Exendin-4 injection (fourth imaging session) of the same islet engrafted into a WT animal exposed to regular chow diet.

**Supplementary video 15. Longitudinal Intravital imaging of islets engrafted into the anterior chamber of the mouse eye, fed with HFHS diet.** *In vivo* calcium imaging at day 8 after vehicle (PBS) injection followed by a Exendin-4 injection (fourth imaging session) of the engrafted into WT animal exposed to regular HFHS diet.

**Supplementary video 16. Longitudinal Intravital imaging of islets engrafted into the anterior chamber of the mouse eye of *Gck*$^{Ki/+}$ animals.** *In vivo* calcium imaging at day 8 after vehicle (PBS) injection followed by a Exendin-4 injection (fourth imaging session) of islets engrafted into Gck$^{Ki/+}$ animal exposed to regular chow diet.

**Supplementary video 17. 3D intravital imaging islets engrafted into the anterior chamber of the mouse eye.** *In vivo* 3D calcium imaging of an islet engrafted into a WT animal exposed to regular chow diet. The calcium waves are initiated at the periphery and propagate across the whole islet.

**Supplementary video 18. 3D intravital imaging islets engrafted into the anterior chamber of the mouse eye.** *In vivo* 3D calcium imaging at day 8 after vehicle (PBS) injection followed by a Exendin-4 injection (fourth imaging session) of islets engrafted into a WT animal exposed to regular chow diet.

**Supplementary video 19. 3D intravital imaging islets engrafted into the anterior chamber of the mouse eye, fed with HFHS diet.** *In vivo* 3D calcium imaging at day 8 after vehicle (PBS) injection followed by a Exendin-4 injection (fourth imaging session) of islets engrafted into a WT animal fed to regular HFHS diet.
